# Microplastics as a novel facilitator for antimicrobial resistance: Effects of concentration, composition, and size on *Escherichia coli* multidrug resistance

**DOI:** 10.1101/2024.08.01.606221

**Authors:** Neila Gross, Johnathan Muhvich, Carly Ching, Bridget Gomez, Evan Horvath, Yanina Nahum, Muhammad H. Zaman

**Affiliations:** Department of Materials Science and Engineering, Boston University, Boston, MA 02215; Department of Biomedical Engineering, Boston University, Boston, MA 02215; Center on Forced Displacement, Boston University, 111 Cummington Mall, Boston, MA, 02215

## Abstract

Microplastics (MPs) have emerged as a significant environmental pollutant with profound implications for public health, particularly as substrates to facilitate bacterial antimicrobial resistance (AMR). Recently, studies have shown that MPs may accommodate microbial communities, chemical contaminants and genetic material containing AMR genes. This study investigated the effects of MP concentration, composition, and size on the development of multidrug resistance in *Escherichia coli*. Specifically, we exposed *E. coli* to varying concentrations of different MP types, including polyethylene (PE), polystyrene (PS), and polypropylene (PP), across a range of sizes (3-10 µm, 10-50 µm, and 500 µm). Results indicated a direct correlation between MP presence and elevated multidrug-resistant (MDR) in *E. coli*.

Notably, MPs exhibited a higher propensity for facilitating resistance than control substrates such as glass, likely due to their hydrophobicity, greater adsorption capacities, and surface chemistries. Furthermore, we observed that co-culture with MPs resulted in biofilm formation. Notably, we found that the bacteria from passaged MPs formed stronger biofilms once the MPs were removed, associated with changes in motility. Thus, we find that MPs also select for cells that are better at forming biofilms, which can lead to recalcitrant infections in the environment and healthcare setting. Our study highlights the immediate need for comprehensive environmental management strategies to mitigate the risk posed by MPs.

**Importance:** Antimicrobial resistance is one of the world’s most pressing global health crises, with an estimated 10 million deaths per year forecasted by 2050. With the pipeline of antibiotics running dry, it is imperative that mitigation strategies understand the mechanisms that drive the genesis of antimicrobial resistance. One emerging dimension of antimicrobial resistance is the environment. This study highlights the relationship between a widespread environmental pollutant, (MPs), and the rise of drug-resistant bacteria. While it is known that MPs facilitate resistance through several modes (biofilm formation, plastic adsorption rates, etc.), this study fills the knowledge gap on how different types of MPs are contributing to antimicrobial resistance.

## Introduction

Global plastic use and mismanaged disposal are significant environmental and public health concerns. Plastic use has increased twenty-fold since 1964, and prevailing estimates suggest global unmanaged trash will reach 155-265 megatons per year in 2060.^1^ Not surprisingly, the detection of (MPs) has significantly increased. MPs are canonically insoluble synthetic particles or polymer matrices with regular or irregular shapes and sizes ranging from the micrometer to millimeter range.^2^ Primary sources of MPs include polyethylene, polypropylene, and polystyrene particles in cosmetic and medicinal products.^3^ Several countries have banned the use of primary MPs within certain industry segments; however, MPs are still generated through the degradation of existing plastics, creating contamination beyond legislation around new product formulations.^3^ These degradation products, characterized as secondary MPs, originate from physical, chemical, and biological processes, resulting in plastic debris fragmentation. Different sources of MPs and their varying surface chemistry also cause them to occur in varying shapes and sizes, such as pellets, fibers, and fragments in environmental samples.^3^ Depending on the conditions plastics are exposed to and the resulting change in surface properties, MPs arise as a unique substrate that has proved to be very difficult to control.^25, 26^

MPs have infiltrated various ecosystems on the planet, ranging from submarine canyons^4^ to the summit of Mount Everest.^5^ Additionally, wastewater has become a significant reservoir for MPs and other anthropological wastes. Despite global awareness, MPs can persist through wastewater treatment plants, disseminating them into surrounding environments. Their persistence can be attributed to their small size, buoyancy, and hydrophobic properties, allowing them to adhere to organic matter and avoid sedimentation.^6^ Consequently, treated wastewater effluents serve as a main source of MP pollution in aquatic environments where they accumulate in sediments and surface waters and interact with the organisms around them.^6^ The repercussions of MPs in wastewater are manifold, especially considering their impacts on human health.

Contemporaneously, increased rates of antimicrobial resistance (AMR)—the ability of microbes to protect themselves against antimicrobials—have been observed in bacterial populations across the globe. AMR can be influenced by a multifaceted network of factors, including the overuse and misuse of antibiotics, poor sanitation and hygiene, and environmental contamination with antibiotic residues in wastewater.^27^ Recent studies show that MPs might also play an important role in the development of AMR.^28, 29, 30^ This is mainly due to their ability to accommodate microbial communities and chemical contaminants and genetic material containing antibiotic-resistant genes (ARGs) through biofilm formation.^19^ In these communities of bacteria that grow together colonizing MP surfaces, ARGs can be transferred to pathogenic bacteria through horizontal gene transfer.^19^ While it has been established that bacteria on the surface of MPs host ARGs, there is limited knowledge on AMR development as a function of MP properties (composition, size, concentration, etc.) as well as their interactions with various antibiotics.^8,11^

Understanding the interplay between AMR and MPs is critical, especially in places with high infection rates and significant plastic waste, such as low-resource settings. The difficulty in treating infectious diseases in these areas, combined with inadequate wastewater treatment— which may result in higher concentrations of MPs—may contribute to the observed increase in AMR cases among vulnerable populations. Therefore, understanding the fundamental interactions between MPs and AMR development is imperative. To date, few studies have looked at the effect of MP size, structure, and other features on the development of drug resistance in the presence of antibiotics. Those that have observed ARGs and increased tolerance to antibiotics.^38,39^ In this study, we investigate the impact of different MP characteristics, including MP concentration, surface composition (between plastics and other materials), size, and surface area, on the *in vitro* development of resistance to ampicillin, ciprofloxacin, doxycycline, and streptomycin in *E. coli*. These antibiotics are broad-spectrum, represent different antibiotic classes, and are readily found in wastewater systems.^12^ More specifically, we probed if MP concentration (no. plastics/µL), size, surface area, and composition (between plastics types) had any impact on (1) bacterial growth alone, (2) antibiotic-specific AMR and (3) multidrug resistance (MDR) between the four antibiotics. We then examined potential mechanisms behind difference in AMR development, specifically studying biofilm formation. Our results identify that MPs play an important and significant role in the development of AMR. Overall, our findings may provide context to wastewater surveillance data and provide insights into waste management and associated disease burdens.

## Methods

### Strains and culture conditions

*E. coli* MG1655 (ATCC 700926) was used in all experiments. All liquid cultures were grown in Lysogeny Broth (Miller, LB) medium under shaking conditions at 180 rpm at 37 °C. First, MPs (Table S.1) were added to the media for 48 hours to allow biofilm surface attachment and full maturity.^40^ Following the initial MP exposure, any additional amendments were added, which included subinhibitory levels of ampicillin (Sigma Aldrich), ciprofloxacin (MP Biomedicals, 86393-32-0), doxycycline (Sigma-Aldrich), streptomycin (Sigma-Aldrich), as indicated below.

### Growth Curve Data

Wild-type *E. coli* MG1655 was cultured at 37 in 4mL of the LB with varying MP concentrations (100, 500, 1000, 2000, 4000, 5000, 10000, and 15000 plastics/μL) in glass culture tubes and sampled at 0, 1, and 24 hours. These samples were plated in triplicate on LB agar to determine the CFU/mL at the time point of interest. Cultures were plated before and after being vortexed, to compensate for cells attached to the MP surface, and CFUs were counted 24 hours after incubation (Fig. 1)

**Figure 1.**
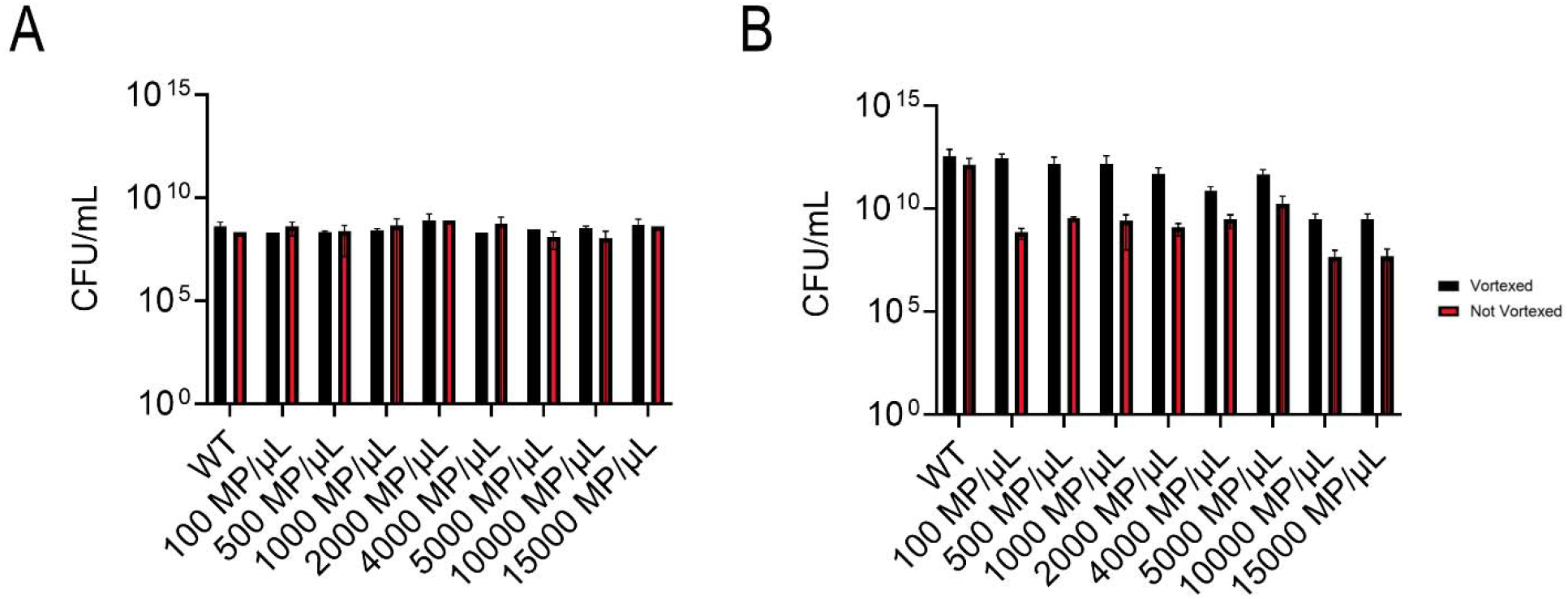
CFU/mL readings after 1 hour of growth (A) and 24 hours of growth (B) in WT and MP concentrations (100, 500, 1000, 2000, 4000, 5000, 10000 and 15000 MP/μL; 10μm diameter polystyrene spheres). Samples that were vortexed are presented in black and non-vortexed samples are depicted in red.

### Antibiotic and MP susceptibility testing

To determine the initial Minimum Inhibitory Concentration (MIC) of bacteria that is used as the Day 0 for the MIC fold change comparison, cells were grown in drug-free media until substantial biofilm growth was detected (∼48 hours) using confocal microscopy (procedure below).

Microparticles (plastics and glass) were vortexed for one minute and spun down to release the biofilm on the surface (Fig. S.1). The supernatant of the vortexed solution was used in a broth microdilution MIC in a 96-well plate using LB media to determine the MIC, effectively testing the MIC of attached cells and effluent cells (Fig. S.2).^34^ The microbroth dilutions begin in the first column with 100μL of an antibiotic and are diluted two-fold into the next column and so on. 10 μL of the samples from the culture are then placed in each well and grown in a static incubator for 24 hours. Following this, the optical density is checked in a spectrophotometer at 600 nm and recorded against a negative control. This was done every 24 hours as shown in Figure S.2.

### MP-antibiotic combination assays

After the initial 48 hour growth period and MIC testing was completed as described above, saturated liquid cultures (cultures that had reach carrying capacity) were passaged once a day into fresh amended media for the antibiotics only or the WT (no antibiotics or MPs)–4 mL of LB broth, 40% of the initial MIC (3.2 μg/mL, 0.0074 μg/mL, 1.6 μg/mL, and 6.4 μg/mL for ampicillin, ciprofloxacin, doxycycline, and streptomycin, respectively) of a singular antibiotic, and their respective growth conditions--via 1:100 dilution until day ten (40μL of the previous day’s culture passaged into 4 mL of the new media). For the MP samples, after each 24-hour exposure period, samples were vortexed to release a portion of the biofilm cells (Fig. S1) to be passaged and tested in MIC plates (Fig. S2B). Not all cells are released from the surface of the biofilm but enough that you can see biofilm in the surrounding media. Then, the MPs were spun down (if they were 10 μm) or allowed enough time to settle (between five and ten minutes) to make sure none were taken out in the following step, in which the supernatant was taken out of the culture tube (40μL saved to passage into the new media and the rest used for the MIC assays), and new media (along with 40μL of the supernatant) was allotted into the *same* culture tube containing the *same* MPs (with less biofilm than pre vortexing) throughout the entire experiment (Fig. S2B). All groups were tested for antibiotic susceptibility every two days, as described above. As a control, cells in drug-free media with the same variable of MPs (concentration, size, composition), cells with no adhesion sites (no MPs or glass spheres) but with drug (40% MIC), as well cells in drug-free and particle free media were also tested (Wild Type (WT)). A schematic of the experiment is displayed in Fig. S.2.

### Confocal Laser Scanning Microscopy (CLSM) image collection and analysis

Visualizing the cells on the surface of the particles was done using confocal microscopy. Using the cultures described in the susceptibility testing section, 500 µm diameter polystyrene spheres were randomly selected out of each culture and dyed using the LIVE/DEAD *Bac*Light Bacterial Viability Kit Protocol, per manufacturer instructions. Briefly, SYTO 9 was used to analyze viable cells, as it can permeate all bacterial cell membranes. In contrast, propidium iodide was used to count dead cells, as it can only enter cells with disrupted membranes. Each plastic was put into a centrifuge tube with 1 mL of sterile water, and three μL of staining solution (1.5 μL of each dye) was injected into the tube. Samples were incubated for 20 minutes at room temperature and protected from light.

Stained samples were imaged with an inverted CLSM (Olympus DSU spinning disk confocal) using a 10X oil immersion objective. Samples individually stained with propidium iodide and SYTO 9 were first analyzed separately to ensure clear signals without overlap. The particles were removed from the dye and suspended in a costar clear round bottom 96 well plate to preserve the MP shape while imaging. After finding the sample and adjusting brightness parameters, z-stacks with an optimized step size were taken for each sample to obtain a 3D visualization of biofilm viability, starting at the base of the spherical particle where growth started and finishing at the top of the sphere where the growth ended.

Z-stacks were analyzed using FIJI.^14^ 3-D renderings were made by taking the .oir file raw data from the microscope and merging the red and green channels. Following the merge, the channels were elucidated by merging the stacks via image -> stacks -> z project with a “Max Intensity” projection type.

### Biofilm formation and Crystal Violet Staining

*E. coli* was inoculated in a 3-to-5-ml culture and grown to stationary phase for 24 hours. The following day, sterile, nontreated 24-well plates (Celltreat ® #229524, Celltreat Scientific Products, Pepperel, MA, USA) were inoculated in 400-μl LB medium per well. The plates were covered and incubated statically at 37° C for 48 hours. After the inoculation period, cells were stained per Merritt et al.’s protocol.^35^ Well optical densities (OD) were then measured in a spectrophotometer at a wavelength of 500 to 600 nm.

### Motility Assay

Following the protocol by (Patridge and Harshey, 2020), 0.3% Agar plates were made and poured to the same thickness (20mL), and let dry for 1 hr.^36^ Using a P20 tip, 5ul of culture was pipetted, inserted into the agar about halfway through, and dispensed. Plates were incubated at 30 C and the diameter of the radial movement (indicated by the visible growth of cells) was then measured using a ruler in centimeters (cm) after 20 hrs.

### Statistical Analysis

The significance of MIC fold changes was determined using an ordinary one-way ANOVA. Before performing the ANOVA, the residuals were tested for normality using the Shapiro-Wilk test, confirming a Gaussian distribution. Each variable was then compared to the mean of a control, which varied depending on the study. Multiple comparisons were corrected with the Dunnett test, with P values adjusted accordingly. Finally, the residuals were tested for homogeneity of variances and potential clustering through the Brown-Forsythe and Bartlett’s tests. The one-way ANOVA was chosen because it allows for comparing mean MIC fold changes across groups under the assumption of normally distributed residuals, which was confirmed here with the Shapiro-Wilk test. Dunnett’s test was applied for post hoc comparisons, as it is well-suited for comparing each group to a control with minimized type I error. Finally, the Brown-Forsythe and Bartlett’s tests were used to assess variance homogeneity, ensuring the robustness of the ANOVA results.

## Results

### MPs impact growth at high concentrations

Eight different concentrations of MPs were grown with *E. coli* for 18 hours to determine whether MPs influence bacterial growth. The MPs used for testing concentration dependence were 10-μm diameter polystyrene spheres at concentrations of 100 MP/µL, 500 MP/µL, 1,000 MP/µL, 2,000 MP/µL, 4,000 MP/µL, 5,000 MP/µL, 10,000 MP/µL, and 15,000 MP/µL. Colony-forming units per mL (CFU/mL) were tested at one hour and 24 hrs after growth in the different concentrations. After one hour of growth, there were no significant differences in growth between the concentrations; however, at 24 hours, two MP solutions (10,000 and 15,000 MP/μL) showed a significantly lower number of CFU/mL compared to the WT (Fig. 1A, 1B). Notably, counts were higher in vortexed samples at 24 hrs, indicating the presence of surface-attached cells on the MPs, which is corroborated by confocal imaging (Fig. S1).

### Exposure to polystyrene MPs increases MDR

We next sought to determine if cells exposed to MPs and subinhibitory levels of a given antibiotic, another common environmental contaminant, had altered patterns of resistance to other classes of antibiotics, rendering the bacteria multidrug-resistant. The four antibiotics we tested were ampicillin (β-lactam antibiotic), ciprofloxacin (fluoroquinolone), doxycycline (tetracycline), and streptomycin (aminoglycoside), which are all commonly found within the environment.^12,15^ First, we measured changes in MIC in cells grown with MPs (500 μm diameter polystyrene spheres due to their ease of passaging) alone, to determine the role of MP on AMR in the absence of antibiotics (Fig. 2). In these susceptibility assays, specifically, we are testing the MIC of effluent containing attached cells released from the MPs upon vortexing (Fig. S1).

**Figure 2.**
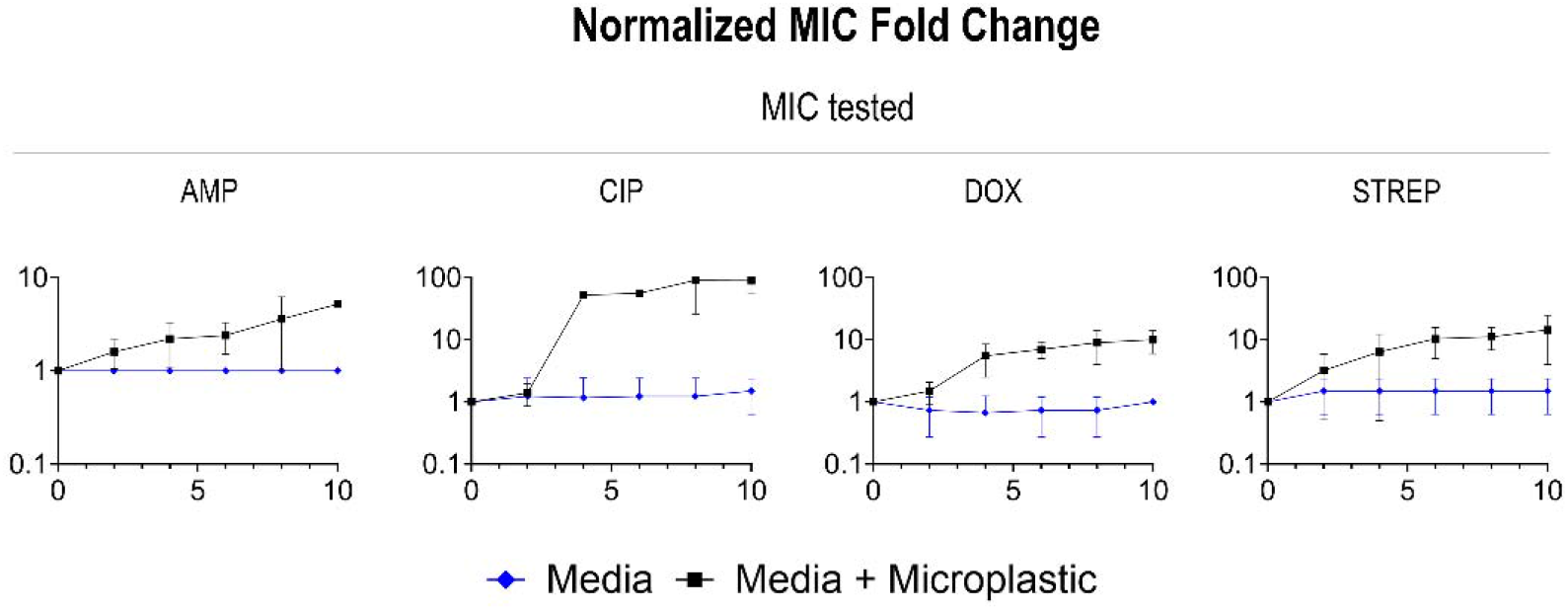
Normalized 10-day time (x-axis) series fold change (y-axis) of 500-μm diameter polystyrene MPs at 40 MP/mL, grown up in LB and then tested for the MIC in all four antibiotics (left to right: ampicillin, ciprofloxacin, doxycycline and streptomycin).

Strikingly, the presence of MPs alone led to increased resistance to all the antibiotics listed above compared to media containing no MPs. While the MIC of bacteria grown in only LB stayed relatively stable, the samples co-cultured with MPs displayed a significantly higher resistance at a faster rate (Fig. 2). Most notably, the cells from the passaged MPs samples had an almost 100- fold higher MIC by day ten compared to day zero for ciprofloxacin, doxycycline and streptomycin (Table S.1). Next, we measured the MIC of cells grown with a single subinhibitory antibiotic with or without MPs (500 μm diameter polystyrene spheres).

Our results suggest that the addition of MPs led to an increase in AMR for nearly all antibiotics. In the first four rows of Fig. 3, we show the fold change of antibiotics relative to WT for bacteria grown in either antibiotics alone or antibiotics and MPs. In each case where bacteria were grown and tested in the same antibiotic, the addition of MPs to antibiotics in the media led to an MIC increase of at least 5 times more compared to cells grown in the antibiotics alone. Interestingly, bacteria grown in ciprofloxacin with MPs generally had higher levels of multidrug resistance (up to 171 times that of the control grown in the antibiotic) (Fig. 3, Table S.1). The other antibiotics with MPs displayed resistance to ciprofloxacin of up to 75-fold higher than the antibiotic control (Table S.1). Additionally, bacteria grown in streptomycin with MPs developed extremely high levels of resistance to ciprofloxacin, doxycycline, and streptomycin (Table S.1).

**Figure 3.**
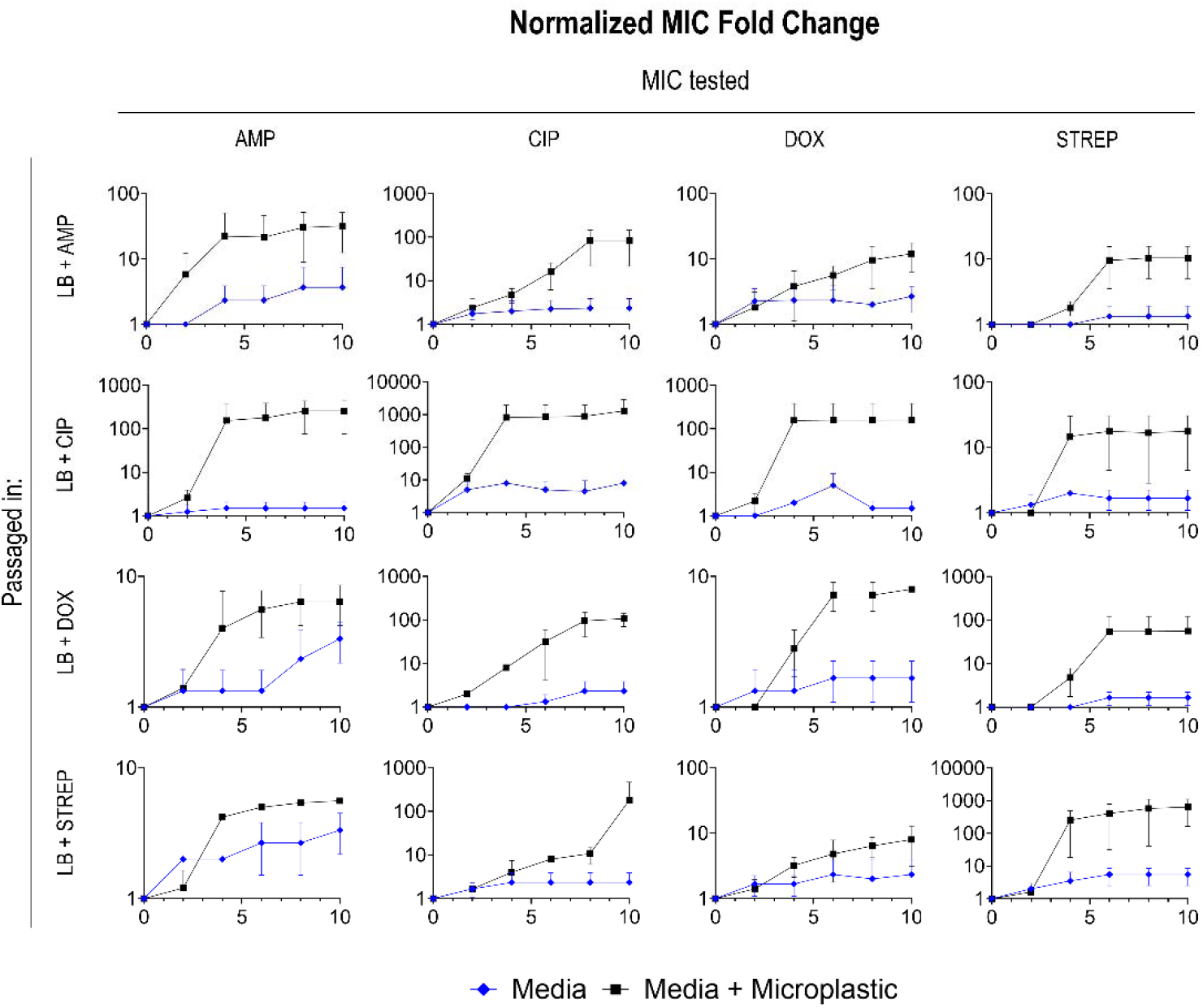
Normalized 10-day time (x-axis) series fold change (y-axis) of 500-μm diameter polystyrene MPs at 40 MP/mL, grown up in one of four antibiotics left to right: ampicillin, ciprofloxacin, doxycycline, and streptomycin and then tested for the MIC in top to bottom: ampicillin, ciprofloxacin, doxycycline and streptomycin

After ten days of exposure to subinhibitory antibiotics, exposure to subinhibitory antibiotics was halted, and bacteria were grown in antibiotic-free media for five days. The MIC was tested daily to determine if resistance was stable. Of the sixteen conditions, 81.25% (13 out of 16) of the bacteria grown in MPs and subinhibitory antibiotics retained the day ten resistance to their respective antibiotic or even gained resistance (Fig. S.3). Conversely, 43.75% (7 out of 16) of the bacteria grown in just MPs retained or grew in resistance to their respective antibiotics.

Surprisingly, only 18.75% (3 out of 16) of the bacteria grown in the subinhibitory antibiotics alone retained resistance. Two of the three conditions that expressed stable resistance was grown in doxycycline (Fig. S.3).

### Exposure to different plastic characteristics (concentration and size) does not affect resistance

Next, we sought to investigate whether different MP characteristics (namely size and concentration) affected antibiotic resistance. For simplicity, we used ciprofloxacin in these experiments as the antibiotic showed the highest resistance changes in the abovementioned studies. Thus, the effect of MP concentration (number of plastics/μL) on the development of ciprofloxacin resistance was investigated under subinhibitory ciprofloxacin conditions. To empirically understand whether the concentration of MPs affected ciprofloxacin resistance development and magnitude of resistance, we exposed the bacteria to subinhibitory levels of ciprofloxacin (7.5e-6 mg/mL, 40% of the initial MIC detailed above) and different concentrations of polystyrene MPs. Resistance was determined in MIC fold changes relative to the control (WT *E. coli* grown without MPs or subinhibitory levels of antibiotics) which were tested every 48 hours.

The concentrations of MPs tested with the 500-μm diameter polystyrene spheres were 40 MP/mL and 100 MP/mL. Conversely, the concentrations tested with the 10-μm polystyrene diameter beads were 1,000 MP/μL, 500 MP/μL, 100 MP/μL, and 10 MP/μL. There was a significant difference in ciprofloxacin MIC fold changes between each concentration of MP and the MP-free counterpart (bacteria with subinhibitory levels of ciprofloxacin but no MPs) as well as the WT (no MPs or antibiotics introduced during growth) (Fig. 4A, C), however, there was no significant difference in changes in ciprofloxacin MIC between the different concentrations of MPs (Fig. 4B, D); however, the absolute MIC values increased up to 3.713 ± 0.66 μg/mL of ciprofloxacin—over three times the Clinical & Laboratory Standards Institute (CLSI) defined 1 μg/mL clinical breakpoint—for the larger diameter beads and 2.508 ± 4 μg/mL of ciprofloxacin for the smaller beads (Table S.3 and S.4).^34^

**Figure 4.**
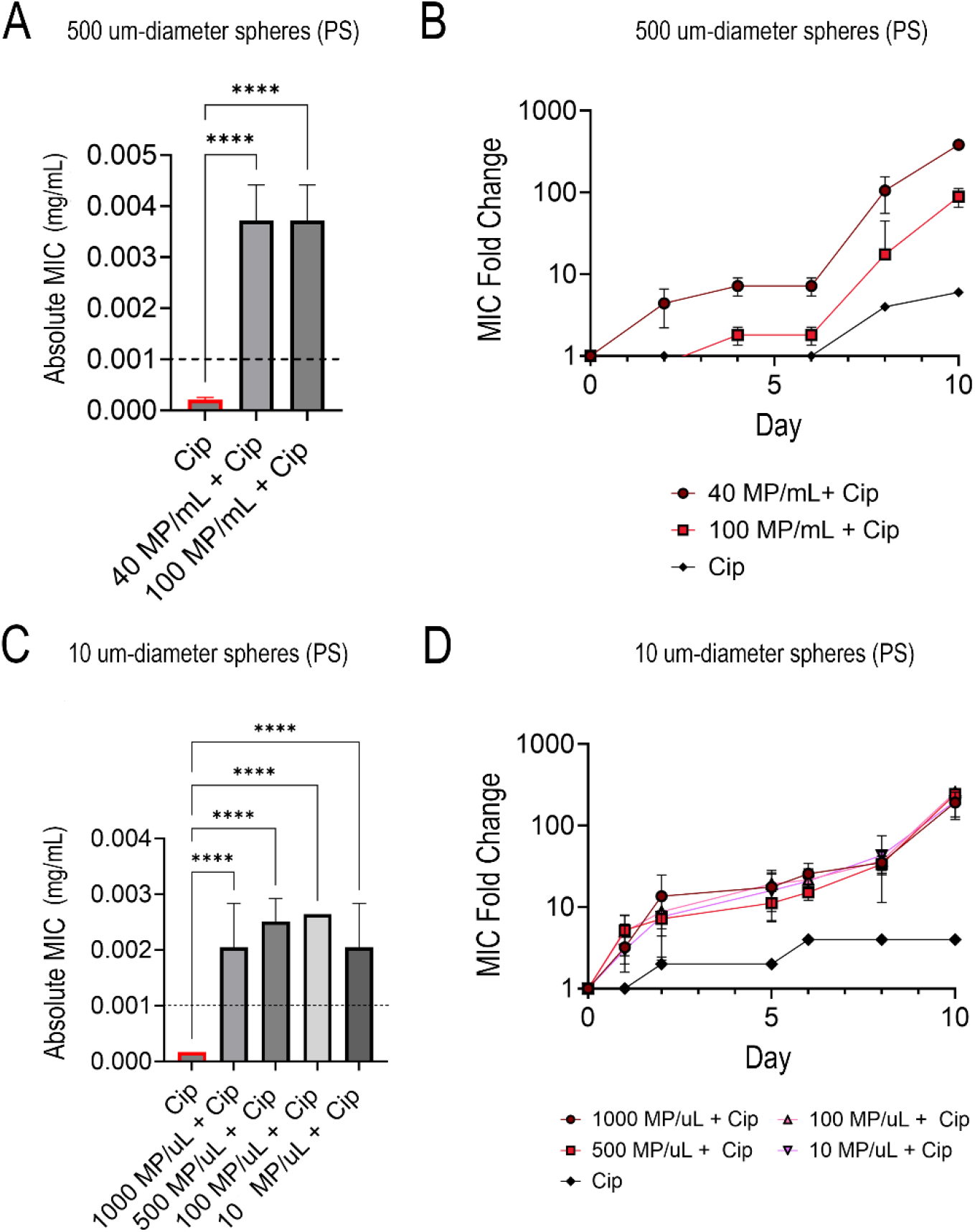
Absolute ciprofloxacin MIC values for 500-μm diameter polystyrene MPs at two different MP concentrations with the dashed line indicating the ciprofloxacin clinical breakpoint concentration at Day 10 of exposure(A), time series fold change of 500-μm diameter polystyrene MPs at different MP concentrations and the subinhibitory antibiotic control (ciprofloxacin) relative to the WT (B), Absolute ciprofloxacin MIC values for 10-μm diameter polystyrene MPs at four different MP concentrations with the dashed line indicating the ciprofloxacin clinical breakpoint concentration at Day 10 of exposure (C)), time series fold change of 10-μm diameter polystyrene MPs at different MP concentrations and the subinhibitory antibiotic control (ciprofloxacin) relative to the WT (D)

MIC variation based on MPs’ size and surface area was investigated following the results above. The sizes of polystyrene MPs investigated were 5 μm (100 MP/μL), 10 μm (100 MP/μL), and 500 μm (40 MP/μL) diameter spheres. All three size variations were exposed to subinhibitory ciprofloxacin along with a control that had no MPs added. Results showed that the difference between the MPs themselves had no significant statistical difference. While the results showed no significant difference between the different surface areas of the beads, the cells grown with MPs and antibiotics again consistently had higher absolute MIC values (above the clinical breakpoint) compared to the antibiotic-only comparators (Fig. 5A, 5B) and the WT.

**Figure 5.**
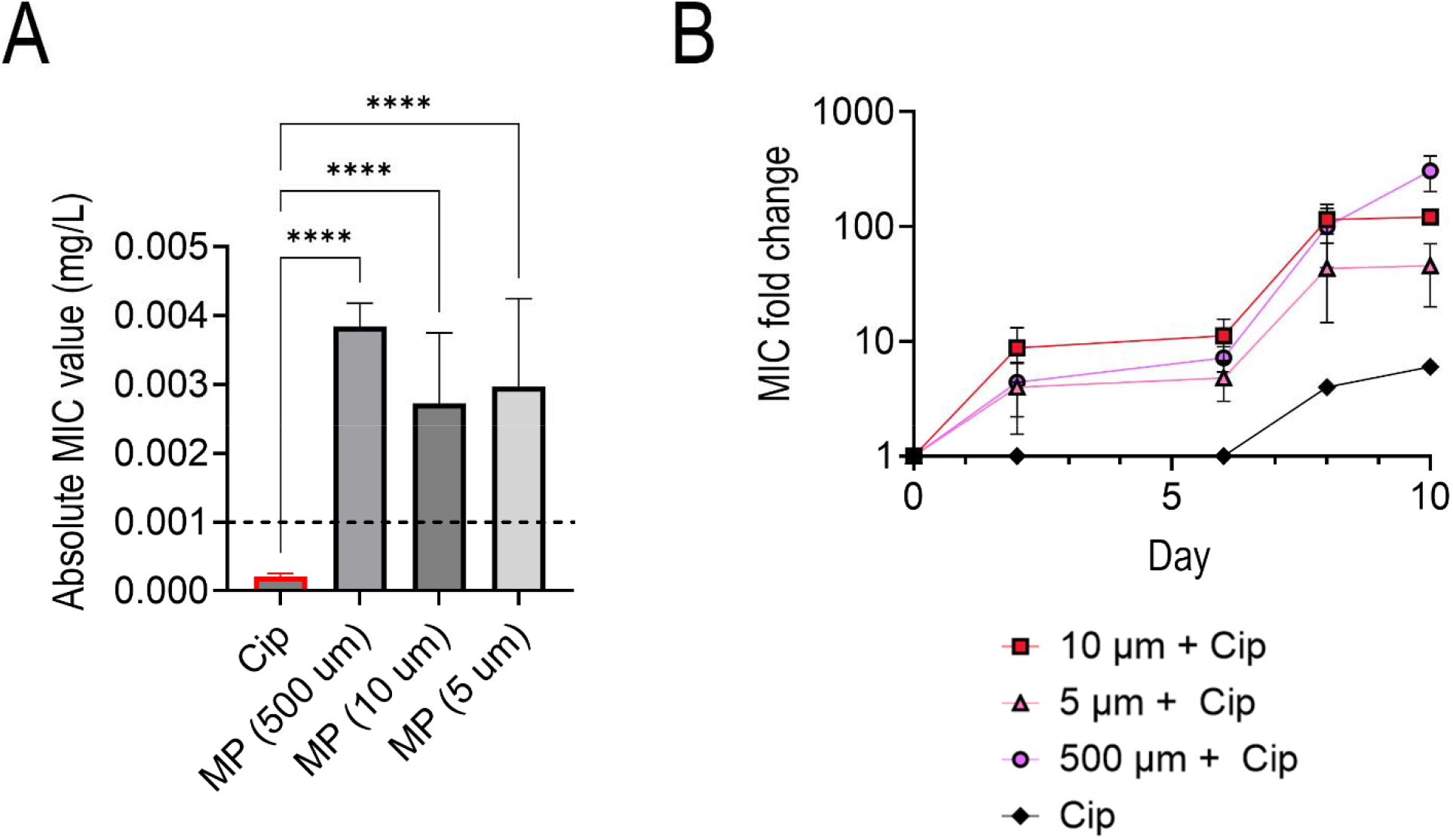
Absolute ciprofloxacin MIC values for various sized polystyrene MPs at 100 MP/μL with the dashed line indicating the ciprofloxacin clinical breakpoint concentration at Day 10 of exposure (A), time series fold change of various sized polystyrene MPs at 100 MP/μL, and the subinhibitory antibiotic control relative to the WT (B)

### Plastic compositions affects antibiotic resistance

Next, we sought to determine if plastic composition affected the development and magnitude of ciprofloxacin resistance (Fig. 6). This experimental design tested and compared the three most common plastic types–polystyrene, polyethylene, and polypropylene.^24^ We found that all three conditions with varying MP compositions had a significantly higher MIC after ten days of exposure than the MP-free control exposed to only subinhibitory levels of ciprofloxacin. The polystyrene samples reached a significantly different absolute MIC value (1.65 ± 0.572 μg/mL) than polyethylene (0.743 ± 0.36 μg/mL), polypropylene (0.528 ± 0.162 μg/mL) and the control of only ciprofloxacin (0.20625 ± 0.04 μg/mL) (Fig. 6, Table S5). Furthermore, the polystyrene MPs facilitated an absolute MIC higher than the ciprofloxacin-*E. coli* clinical breakpoint (1 μg/mL). Interestingly, the rates of AMR growth in all three plastic samples were similar in terms of fold change (Fig. 6B).

**Figure 6.**
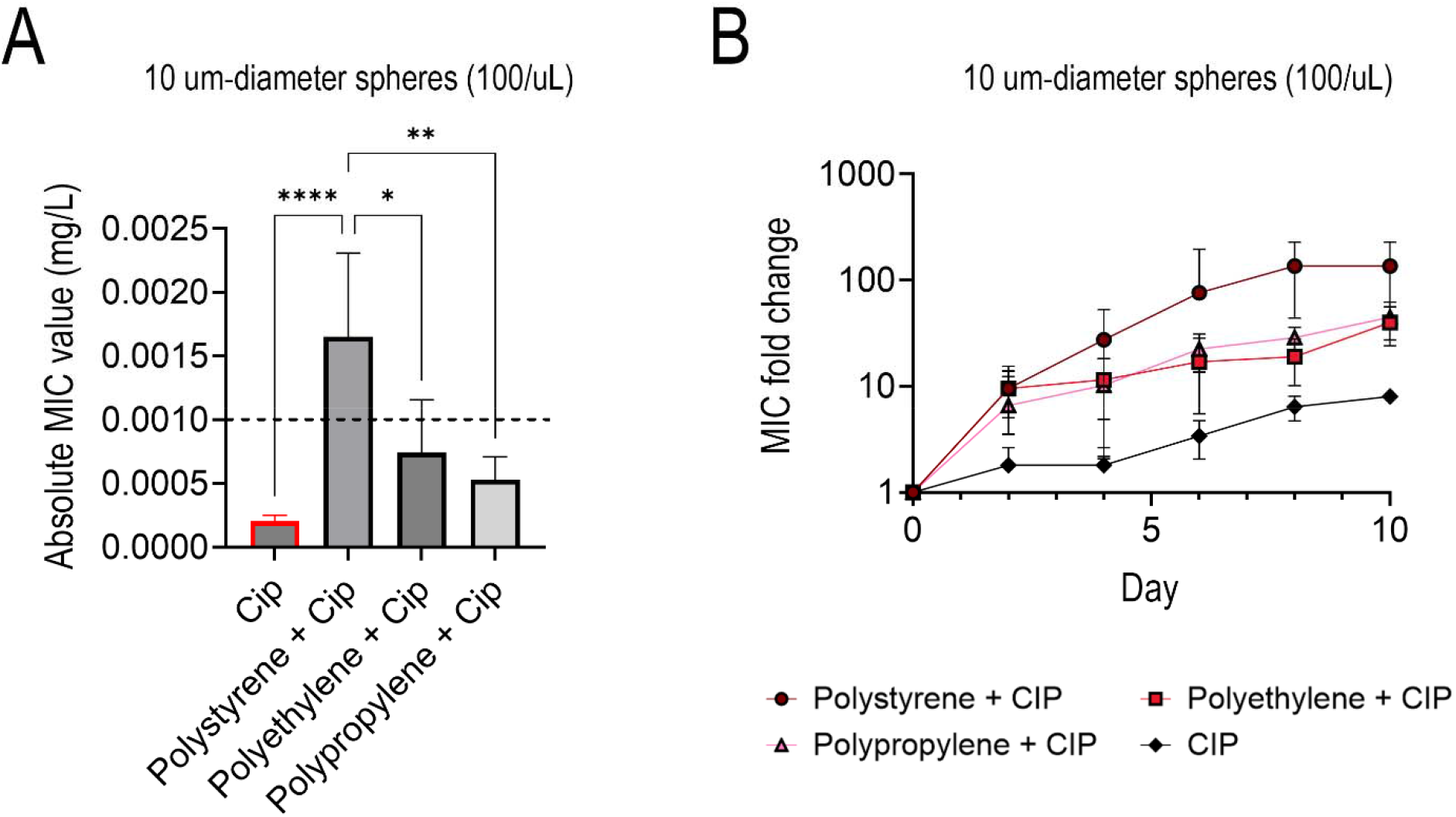
Absolute ciprofloxacin MIC values for 10-μm diameter polystyrene, polyethylene, and polypropylene MPs at 100 MP/μL with the dashed line indicating the ciprofloxacin clinical breakpoint concentration at Day 10 of exposure (A), time series fold change of 10-μm diameter polystyrene, polyethylene, polypropylene MPs at 100 MP/μL, and the subinhibitory antibiotic control relative to the WT (B)

### Increased resistance and biofilm on polystyrene compared to glass

We also studied the difference in resistance development between polystyrene MP and glass spheres (500-μm diameter spheres at the same concentration of 40 MP/μL) to determine if plastic substrates had a specific effect or if any small particles would lead to an increase in AMR (Fig. 7). Similar to previous experiments, the microparticles were exposed to subinhibitory levels of ciprofloxacin and compared to a control, which only had the subinhibitory ciprofloxacin and no glass or plastic particles. The polystyrene plastic spheres facilitated a statistically higher absolute MIC value by the end of the ten-day study from its glass counterpart and antibiotic control (Fig. 7A). Like Figure 6, the fold changes of the attachment surfaces were not statistically different (Fig. 7B). This indicates that the rate of AMR growth is similar in terms of fold change.

**Figure 7.**
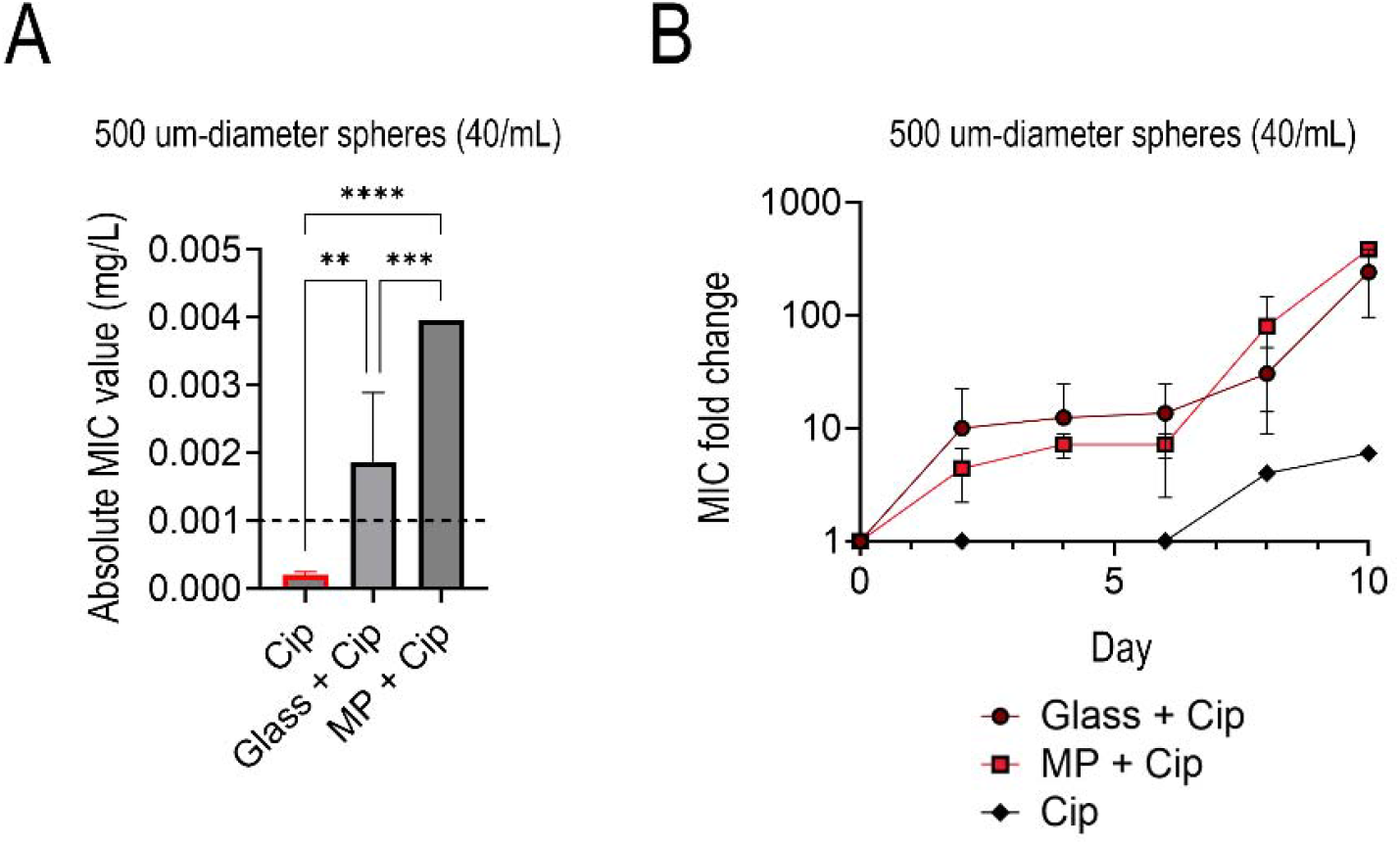
Absolute ciprofloxacin MIC values for 500-μm diameter polystyrene and glass spheres at 40 MP/mL at Day 10 of exposure (A), with the dashed line indicating the ciprofloxacin clinical breakpoint concentration. Time series fold change of the 500-μm (B) glass, polystyrene, and the subinhibitory antibiotic control relative to the WT.

To elucidate the mechanism behind the high levels of resistance found in the samples containing the MPs (particularly polystyrene, Fig. 2 and 3) and the discrepancies behind the resistance profiles of the particles of different compositions, we wanted to visualize the surface of the particles. We did this qualitatively by using confocal microscopy. The glass and polystyrene spheres were dyed on day 10 of subinhibitory ciprofloxacin exposure and compared to each other (Fig. 8) using a live dead stain. Green pixels indicate viable growth (e.g., live cells). In contrast, red indicates nonviable growth (dead cells). The glass (Fig. 8A) depicts two spheres, each having a significantly decreased amount of growth on the surface of the sphere, and most of the growth exhibited was colonized by dead cells. Conversely, the polystyrene condition had mostly live cells, and bacteria colonized the whole surface.

**Figure 8.**
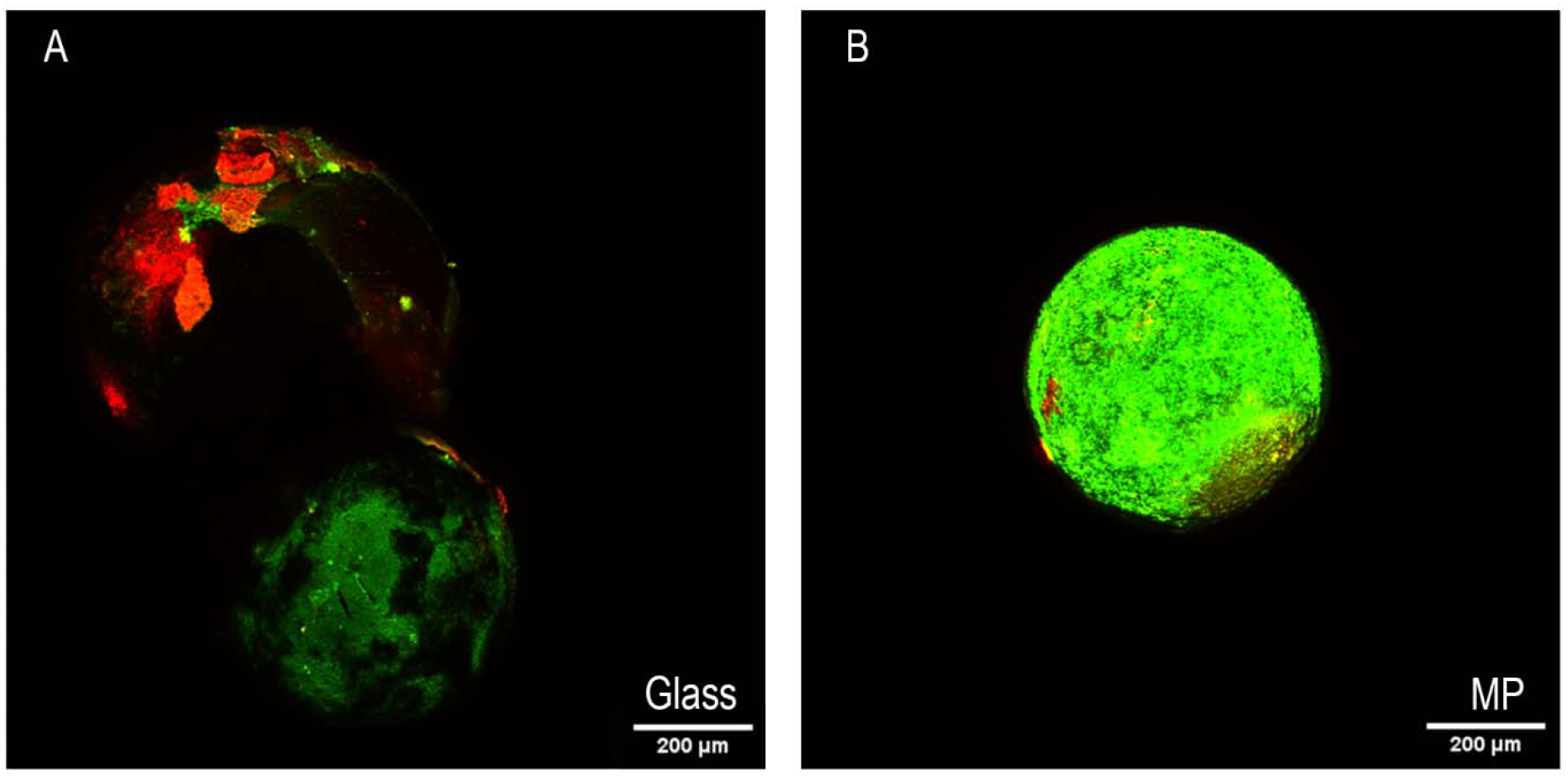
CLSM image of 500-μm diameter glass (left) and polystyrene (right) spheres at day 10 of subinhibitory ciprofloxacin exposure. Green pixels indicate live cells, while red indicate dead cells.

### Exposure to MPs select for increased biofilm

Following the visualization of robust biofilms on the surfaces of the polystyrene MPs, we sought to understand the role of biofilm further. Specifically, we investigated whether cells passaged with MPs (Fig. 2 and 3) formed more robust biofilms in the absence of MPs (i.e. does the presence of MPs select for better biofilm formers). Crystal violet staining can detect biofilm formation as it binds to bacterial cells and the extracellular matrix, making biofilms both visible and quantifiable. The bacteria grown with and without MPs along with various antibiotics and controls for 10 days (Fig. 2 and 3) were purified of MPs and trace antibiotics and grown in a 24- well polystyrene dish for 48 hours, and biofilm (surface attachment) was quantified using 0.1% crystal violet solution (Fig. 9).

**Figure 9.**
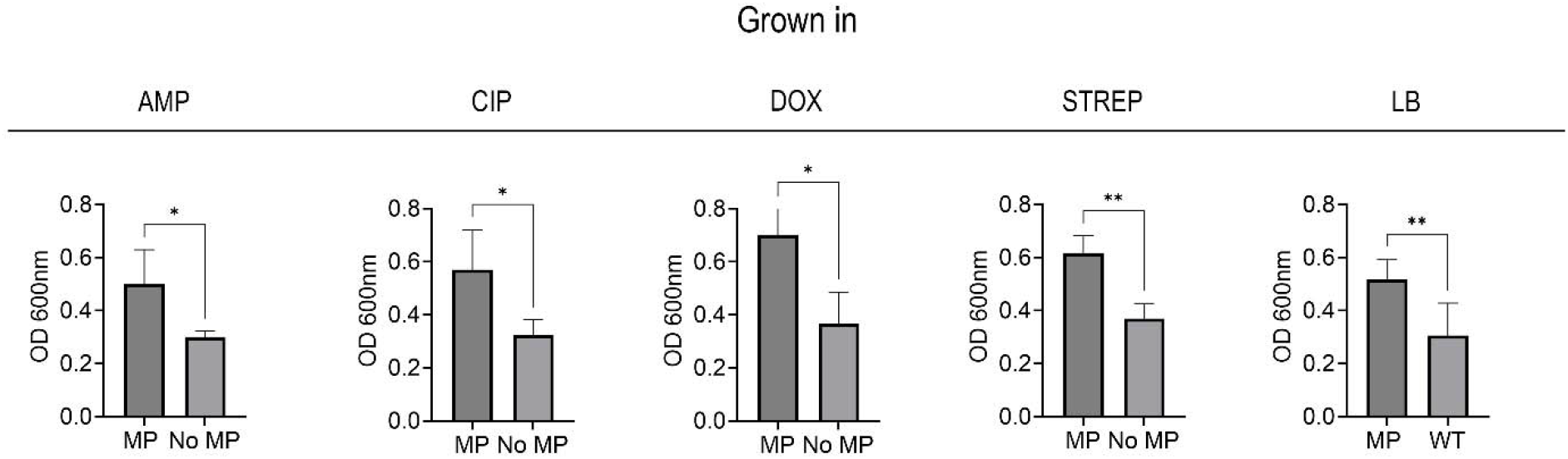
OD readings of crystal violet staining for E. coli grown with or without MPs in media containing subinhibitory levels of four antibiotics left to right: ampicillin, ciprofloxacin, doxycycline, or streptomycin or no antibiotics (LB).

All samples grown with MPs had significantly more biofilm growth than those grown without MPs, indicated by their higher OD readings and increase in stained surface-attached cells (Fig. 9, Fig. S.4). Bacterial motility is a broad mechanism involved in biofilm formation; specifically, impaired motility is associated with increased biofilm due to several interconnected factors, mainly related to regulatory shifts, changes in surface properties, and environmental sensing mechanisms.^37^ Thus, swimming motility was also assayed using the bacteria grown with and without MPs along with various antibiotics and controls for 10 days (Fig.2 and 3) on 0.3% soft agar. Cells not grown with MPs had significantly larger diameters than those grown with MPs, indicating that the bacteria exposed to MPs had impaired motility (Fig. 10, Fig. S.5).

**Figure 10.**
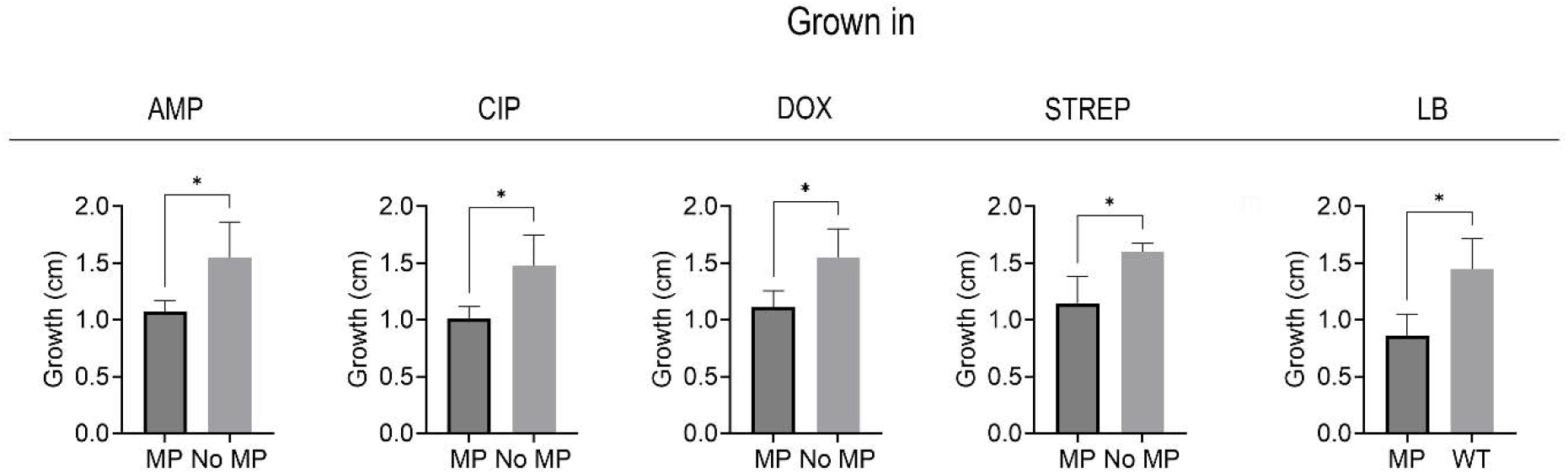
Motility in centimeters (cm) of E. coli grown with or without MPs in media containing subinhibitory levels of four antibiotics left to right: ampicillin, ciprofloxacin, doxycycline, or streptomycin or no antibiotics (LB).

## Discussion

MPs are significant environmental pollutants with critical implications for public health, particularly in the context of increasing AMR. Understanding how MPs affect the development of AMR is a crucial step in better understanding how environmental factors shape AMR in bacteria. This will only become more important as both MPs and AMR become more prevalent. Our study showed that MPs significantly impact the AMR rate of development and its magnitude. More specifically, MPs facilitated post-breakpoint multidrug resistance to four distinct families of antibiotics (Fig. 2A, 2B). Breakpoints are an integral part of modern microbiology practice and define susceptibility and resistance to antibacterials in the clinical setting (e.g., related to human health).^16^

Polystyrene, in particular, had the most significant impact on resistance development, which was surprising given its relative hydrophilicity compared to polyethylene and polypropylene (Fig. 6). Previous studies have shown that bacteria have an affinity for plastic due to their high hydrophobicity and oxidation, which can lead to easy adhesion.^31, 32^ However, it is essential to note that studies have shown that *E. coli* prefers hydrophilic surfaces over hydrophobic ones.^17^ Of the three plastics used in this study, polystyrene is the most hydrophilic, while polypropylene is the most hydrophobic, so these findings are in line with *E. coli’s* preference to hydrophilic surfaces. With this in mind, we expected glass beads of similar diameter to have a higher adhesion rate and, therefore, a higher resistance to the tested antimicrobials. This, however, was not the case, as polystyrene had a higher absolute MIC value and greater MIC fold change over the glass condition. This indicates that plastics may be a unique substrate for bacteria to develop and maintain resistance to.

While the complete mechanism is not yet known for AMR on MPs, the current prevailing theory indicates that biofilm formation upon the plastics allows for higher resistance rates.^29^ As depicted in the CLSM images in Fig. 8, we observed a dense biofilm that spans the entire surface of the MPs, compared to the glass substrate, which has uneven clusters of cells on the surface, with a large portion of them nonviable post-subinhibitory antibiotic exposure. This would help explain the differences in MIC shown in Fig. 7. We investigated the effects the cells had post-MP exposure to see if they had a propensity for forming biofilms over their counterparts grown in the same media without MPs. Qualitatively, confocal imaging found that not only did the bacteria grown with MPs develop biofilms, but they grew biofilms with more biomass on the plastics than glass surfaces (Fig. 8). Notably, cells grown in subinhibitory levels of doxycycline and MPs had the highest biofilm growth. Doxycycline was the only bacteriostatic antibiotic—an antibiotic that attacks cell reproduction rather than the cell itself—used in this study.

Doxycycline is known to target the 30S ribosomal subunit and inhibit protein synthesis, which can trigger stress pathways that upregulate biofilm-associated genes and extra polymeric substance production.^41^ We next examined bacterial motility to investigate further the bacteria’s ability to grow biofilms. In these experiments, we found that samples grown with MPs had impaired motility, which can influence biofilm formation (Fig. 10).^37^ Overall, these data suggest that the presence of MP not only presents a surface for biofilm formation but also selects for cells that are better at biofilm formation. We hypothesize that, especially in the presence of low levels of antibiotics, there is a combinatorial effect between selection for antibiotic resistance target genes and biofilm formation that leads to high levels of resistance and recalcitrance.

Further research is needed to investigate the cellular-level interactions and material properties of plastics that contribute to such high resistance rates.

It is well known that biofilms play a crucial role in the spread of AMR. Bacteria within biofilms produce persister cells that are metabolically inert, which is one mechanism for evading antibiotics.^22^ These cells can survive even in high concentrations of antibiotics.^22^ Furthermore, current research suggests that biofilms act as refugia of MDR plasmids by retaining them, even in the absence of antibiotics.^23^. This would support our results with respect to the MP properties we investigated. First, MP concentration was not factored into different resistance rates. Instead, size and composition had more of an impact on the rate of resistance development and magnitude of resistance. A larger-sized MP would have a larger biofilm and, therefore, a greater capacity to develop resistance. The plastic composition can also play a role in both bacterial attachment and biofilm growth. MPs are known to serve as electron donors for bacterial biofilms to feed on, inducing a faster or easier attachment of the bacteria to the surface and promoting bacterial growth and colonization.^31, 35^ This may explain the higher concentration of biofilm on polystyrene MP compared to glass (Fig. 7), which in turn would explain polystyrene’s larger resistance load.

The proposed mechanisms of biofilms and selective pressures are assumed to have accumulated overtime and compounded off of each other, creating high resistance levels as the time trial went on. We believe that the potential ramifications of high-level multidrug-resistant bacteria facilitated by the addition of MPs are significant. Moreover, we found bacteria exhibiting this behavior within ten days of subinhibitory antibiotics and MP exposure. The rate of AMR development and the surpassing of clinical breakpoints in both single and multidrug tests highlight a need to monitor MPs and antibiotic levels in the environment. This is especially true in areas with inadequate waste disposal and substandard public health infrastructure, such as low-and middle-income countries (LMICs) and vulnerable populations.^18^ Future studies in this area should focus on the disparities of wastewater treatment in LMICs and high-income countries and how the different environmental factors shape AMR development. Additionally, wastewater and environmental monitoring should also include the presence of MPs, as they have the potential to exacerbate AMR outbreaks. Our work can inform the ongoing development of AMR surveillance strategies, helping to predict and prevent future outbreaks.

## Acknowledgments

Thank you to Matt Lim for guidance at the confocal microscope and Boston University’s shared CORE facility for the usage of their instruments.

## CRediT Statement

NG: Conceptualization, Methodology, Investigation, Writing-Original Draft, Writing- Review and Editing

JM: Investigation, Writing-Review and Editing

CC: Conceptualization, Writing-Review and Editing

BG: Investigation, Writing-Review and Editing

EH: Writing-Review and Editing

YN: Writing- Review and Editing

MHZ: Writing-Review and Editing

## Data Sharing

All data used for this study has been included in the manuscript or supplementary material.

