## Supplemental figures for "Microplastics as a novel facilitator for antimicrobial resistance: Effects of concentration, composition, and size on *Escherichia coli* multidrug resistance"

Supplementary Information

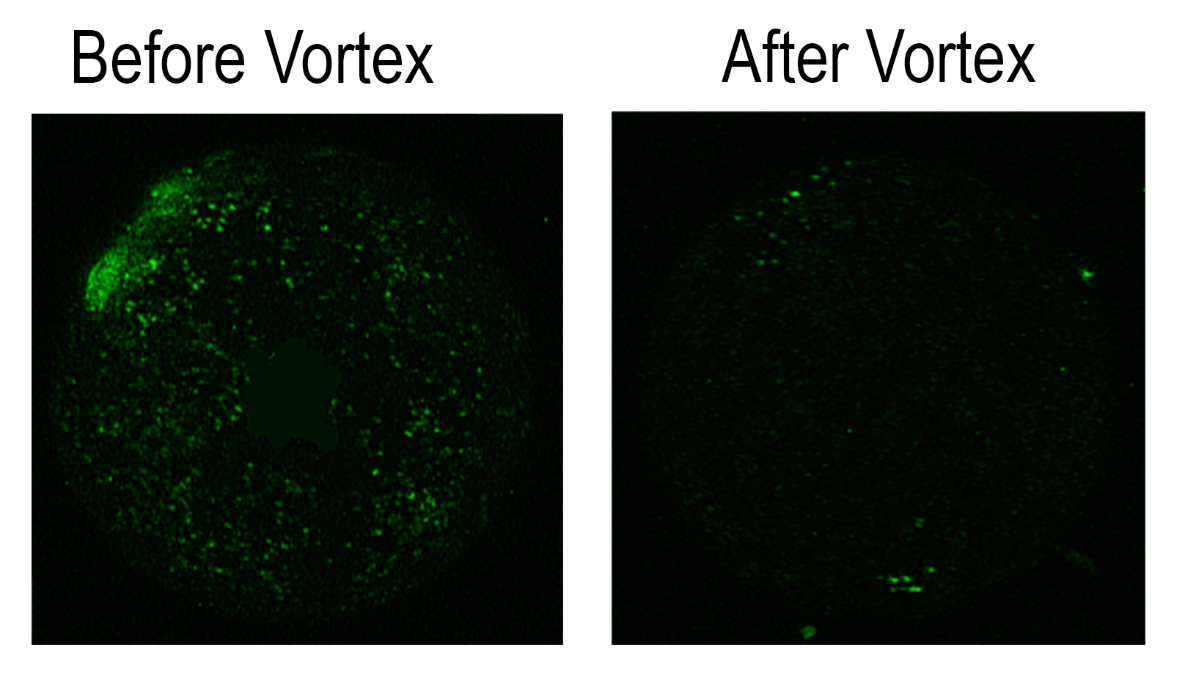

***Figure S.1.*** *MP before and after the sample was vortexed for one minute. Green cells indicate live E. coli cells attached to the surface of the MP.*

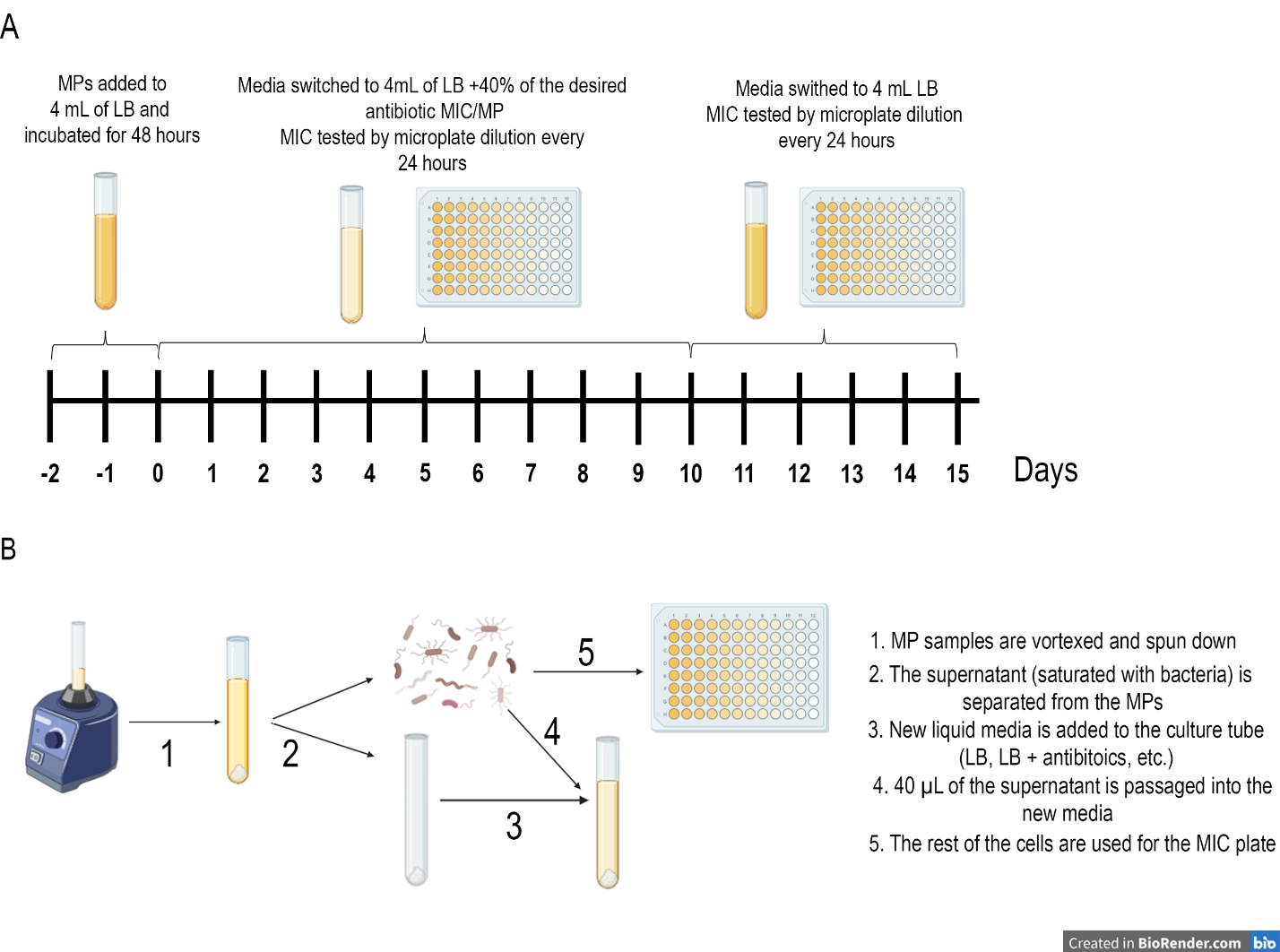
***Figure S.2.*** *Experimental schematic: Cells were pre-incubated with or without microplastics for two days before subinhibitory antibiotic exposure. Starting from day zero, the minimum inhibitory concentration (MIC) was measured in a standard microdilution plate with the antibiotic of interest before returning to being passaged in just LB for days 10-15 (A), Detailed passaging schematic for microplastic samples (B).*

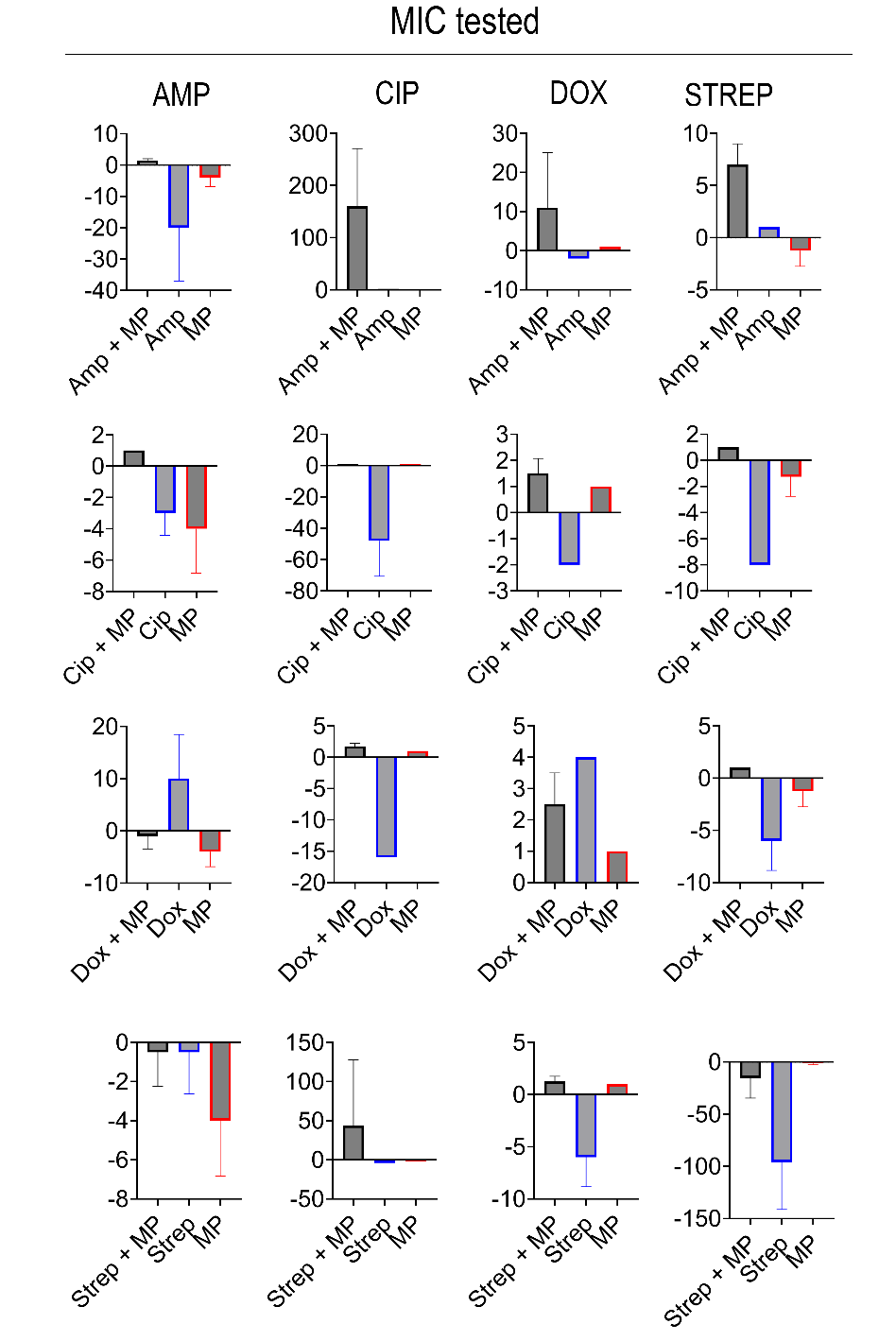

***Figure S3.*** *Five-day resistance stability was measured in fold change (y-axis) relative to day 10 of the MDR study above. The bacteria grown with subinhibitory antibiotics + MP (black), bacteria + subinhibitory antibiotics (blue), and bacteria + MPs (red) (x-axis) are shown on day 15 of exposure to LB only. Negative values indicate MIC fold change* lost*, while positive values indicate MIC fold change increase.*

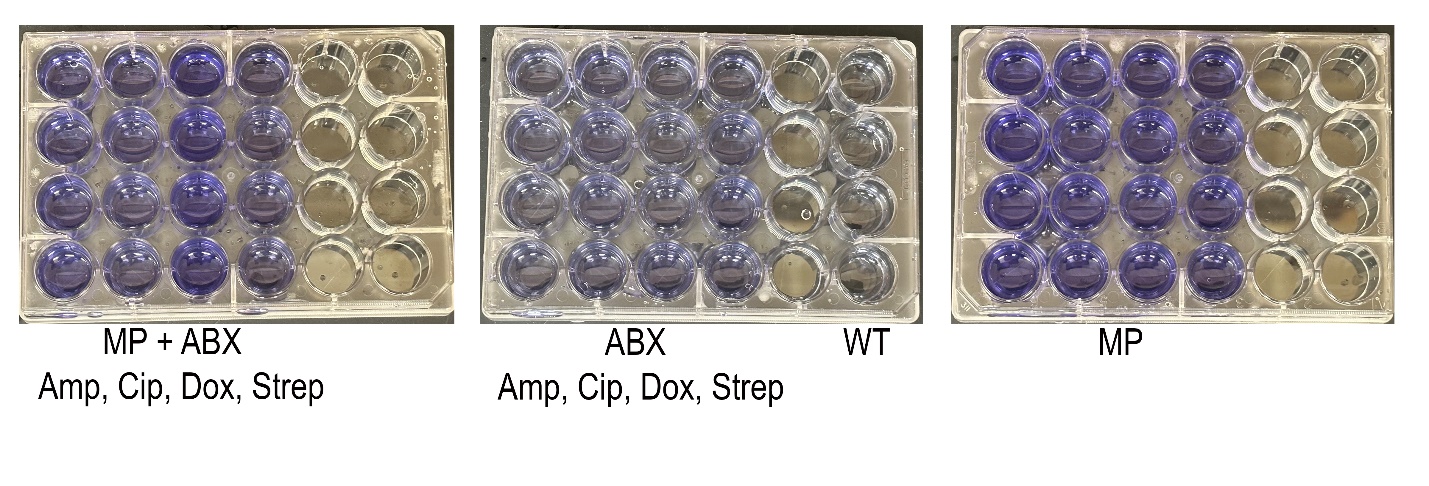

***Figure S.4****. 0.1% Crystal Violet stains on bacterial samples post 10-day exposure to various media (subinhibitory ampicillin, ciprofloxacin, doxycycline, streptomycin, or LB) and with or without MPs. Darker purple indicates more biofilm growth.*

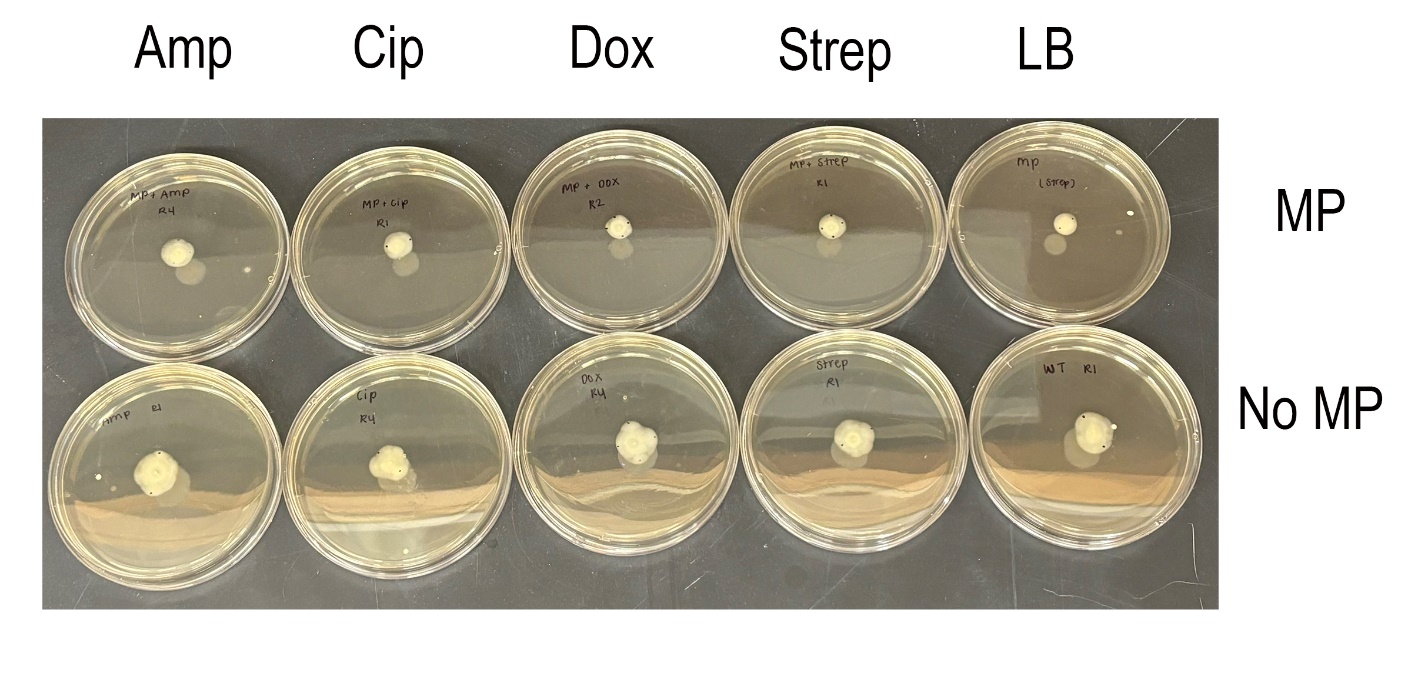

***Figure S.5****. Sample of soft agar plates to determine motility of the bacterial* *samples post 10-day exposure to various media (subinhibitory ampicillin, ciprofloxacin, doxycycline, streptomycin, or LB) and with or without MPs. Larger cell growth radius indicates higher motility.*

***Table S.1*** *Microplastics and their vendors*

| *Composition* | *Size* | *Vendor* |
| --- | --- | --- |
| Polystyrene spheres | 500-600 μm | Polysciences, cat. 21392-1 |
| Polystyrene spheres | 10 μm | Alpha Chemistry, cat ACM9003536-5 |
| Polyethylene spheres | 10 μm | Alpha Chemistry, cat. ACM9002884-4 |
| Polypropylene spheres | 10 μm | Alpha Chemistry, cat. ACM9010791-5 |
| Silica (glass) spheres | 500 μm | Benchmark Scientific, cat. D1131-05 |

***Table S.2*** *Absolute MIC values (mg/mL) and post-antibiotic exposure fold change for multidrug resistance (MDR) of bacteria + 500-μm polystyrene MPs (40/mL) + antibiotics grown in one antibiotic and tested in four antibiotics, (ii.) bacteria + antibiotics grown in and (iii.) bacteria + 500-μm polystyrene MPs (40/mL) grown in antibiotics and tested in four antibiotics at Day 10 of exposure.*

|  | 500 μm diameter MP | Absolute MIC (μg/mL) | Fold Change (unitless) |
| --- | --- | --- | --- |
|  | Tested in: | *Grown in: Ampicillin + MP* | |
|  | Amp | 100 ± 36.7 | 32 ± 17.5 |
|  | Cip | 0.4125 ± 0.147 | 83.2 ± 54.87 |
|  | Dox | 1.563 ± 1.7 | 12 ± 5.06 |
|  | Strep | 7.813 ± 2.296 | 10.4 ± 4.8 |
|  | Tested in: | *Grown in: LB + Ciprofloxacin + MP* | |
|  | Amp | 6.25 ± 2.5 | 256.8 ± 160.6 |
|  | Cip | 3.96 ± 1.833* | 1305.6 ± 1442.26 |
|  | Dox | 1.563 ± 0.7 | 160 ± 200 |
|  | Strep | 15.625 ± 1.53 | 17.6 ± 11.76 |
| Tested in: | | *Grown in: LB + Doxycycline + MP* | |
| Amp | | 10 ± 7.5 | 6.4 ± 1.96 |
| Cip | | .33 ± 0.808 | 106.67 ± 30.17 |
| Dox | | 2.5 ± 0.7 | 8 ± 0 |
| Strep | | 13.75 ± 1.5 | 56 ± 58.8 |
| Tested in: | | *Grown in: LB + Streptomycin + MP* | |
| Amp | | 5.625 ± 1.25 | 5.6 ± 5.276 |
| Cip | | .825 ± 1.0314 | 181.33 ± 233.82 |
| Dox | | 1.875 ± 0.625 | 8 ± 4.38 |
| Strep | | 100 ± 244.9 | 640 ± 426.45 |
| Tested in: | | *Grown in: LB + Ampicillin* | |
| Amp | | 14.06 ± 10.9 | 1.5 ± 0.5 |
| Cip | | 0.0425 ± 0.03996 | 2.33 ± 1.25 |
| Dox | | 1.953 ± 1.17 | 2.67 ± 0.94 |
| Strep | | 5.078 ± 1.17 | 1.33 ± 0.47 |
| Tested in: | | *Grown in: LB + Ciprofloxacin* | |
| Amp | | 3.907 ± 2.344 | 1.5 ± 0.5 |
| Cip | | 0.0928 ± 0.072188 | 8 ± 0 |
| Dox | | 1.758 ± 1.367 | 1.5 ± 0.5 |
| Strep | | 4.102 ± 2.149 | 2 ± 0 |
| Tested in: | | *Grown in: LB+ Doxycycline* | |
| Amp | | 4.688 ± 1.563 | 3.33 ± 0.943 |
| Cip | | 0.0219 ± 0.019336 | 2.33 ± 1.25 |
| Dox | | 1.172 ± 0.391 | 1.67 ± 0.47 |
| Strep | | 2.539 ± 0.586 | 1.33 ± 0.47 |
| Tested in: | | *Grown in: LB + Streptomycin* | |
| Amp | | 3.907 ± 2.344 | 3.33 ± 0.9428 |
| Cip | | 0.0219 ± 0.019336 | 2.33 ± 1.25 |
| Dox | | 0.977 ± 0.586 | 2.5 ± 1.5 |
| Strep | | 115.625 ± 84.375 | 5.5 ± 1.598 |
| Tested in: | | *Grown in: LB + MP* | |
|  | Amp | 10.938 ± 8.119 | 6 ± 5.83 |
|  | Cip | .33 ± 0 | 42.46 ± 31.35 |
|  | Dox | 3.125 ± 0 | 10 ± 3.46 |
|  | Strep | 31.25 ± 13.258 | 16 ± 9.798 |

***Table S.3*** *Absolute MIC values (μg/mL) and post-antibiotic exposure fold change for MP concentration study with 10-μm polystyrene MPs at Day 10 of exposure.*

| 10 μm diameter PS MP | Absolute MIC (μg/mL) | Fold Change (unitless) |
| --- | --- | --- |
| 1000 MP/uL | 2.046 ± 0.7* | 192 ± 57.2 |
| 500 MP/uL | 2.508 ± 4* | 243.2 ± 25.6 |
| 100 MP/uL | 2.64 ± 0* | 256 ± 0 |
| 10 MP/uL | 2.046 ± 0.7* | 198.4 ± 71.26 |
| Control (ciprofloxacin) | 0.165 ± 0 | 4 ± 0 |

** indicates past ciprofloxacin breakpoint of 1 μg/mL*

***Table S.4*** *Absolute MIC values (μg/mL) and post-antibiotic exposure fold change for MP concentration study with 500-μm diameter polystyrene MPs at Day 10 of exposure.*

| 500 μm diameter PS MP | Absolute MIC (μg/mL) | Fold Change (unitless) |
| --- | --- | --- |
| 40 MP/mL | 3.713 ± 0.66* | 346.8 ± 74.4 |
| 100 MP/mL | 3.7125 ± 0.65* | 88.8 ± 20.4 |
| Control (ciprofloxacin) | 0.165 ± 0 | 4 ± 0 |

** indicates past ciprofloxacin breakpoint of 1 μg/mL*

***Table S.5*** *Absolute MIC values (μg/mL) and post-antibiotic exposure fold change for plastic composition study 10-μm diameter MP spheres at 100 MP/μL concentration at Day 10 of exposure.*

| 10 μm diameter MP | Absolute MIC (μg/mL) | Fold Change (unitless) |
| --- | --- | --- |
| Polystyrene | 1.65 ± 0.572* | 136 ± 79.6 |
| Polyethylene | 0.743 ± 0.36 | 40 ± 13.86 |
| Polypropylene | 0.528 ± 0.162 | 44.8 ± 16.68 |
| Control (ciprofloxacin) | 0.20625 ± 0.04 | 8 ± 0 |

** indicates past ciprofloxacin breakpoint of 1 μg/mL*

***Table S.6*** *Absolute MIC values (μg/mL) and post-antibiotic exposure fold change for microparticle composition study of various sized spheres and surface areas at Day 10 of exposure.*

| 3-5 μm or 500 μm diameter | Absolute MIC (μg/mL) | Fold Change (unitless) |
| --- | --- | --- |
| 500 μm Glass | 1.86 ± 0.89* | 271.2 ± 149.4 |
| 500 μm MP | 3.84 ± 0.317* | 168 ± 128.8 |
| Control (ciprofloxacin) | 0.20625 ± 0.04 | 7.6 ± 0.8 |

** indicates past ciprofloxacin breakpoint of 1 μg/mL*
